# Deciphering regulatory DNA sequences and noncoding genetic variants using neural network models of massively parallel reporter assays

**DOI:** 10.1101/393926

**Authors:** Rajiv Movva, Peyton Greenside, Georgi K. Marinov, Surag Nair, Avanti Shrikumar, Anshul Kundaje

**Affiliations:** The Harker School, San Jose, CA, USA; Department of Genetics, Stanford University, Stanford, CA, USA; Biomedical Informatics Training Program, Stanford University, Stanford, CA, USA; Department of Computer Science, Stanford University, Stanford, CA, USA

## Abstract

The relationship between noncoding DNA sequence and gene expression is not well-understood. Massively parallel reporter assays (MPRAs), which quantify the regulatory activity of large libraries of DNA sequences in parallel, are a powerful approach to characterize this relationship. We present MPRA-DragoNN, a convolutional neural network (CNN)-based framework to predict and interpret the regulatory activity of DNA sequences as measured by MPRAs. While our method is generally applicable to a variety of MPRA designs, here we trained our model on the Sharpr-MPRA dataset that measures the activity of ~500,000 constructs tiling 15,720 regulatory regions in human K562 and HepG2 cell lines. MPRA-DragoNN predictions were moderately correlated (Spearman *ρ* = 0.28) with measured activity and were within range of replicate concordance of the assay. State-of-the-art model interpretation methods revealed high-resolution predictive regulatory sequence features that overlapped transcription factor (TF) binding motifs. We used the model to investigate the cell type and chromatin state preferences of predictive TF motifs. We explored the ability of our model to predict the allelic effects of regulatory variants in an independent MPRA experiment and fine map putative functional SNPs in loci associated with lipid traits. Our results suggest that interpretable deep learning models trained on MPRA data have the potential to reveal meaningful patterns in regulatory DNA sequences and prioritize regulatory genetic variants, especially as larger, higher-quality datasets are produced.

## 1. Introduction

Changes in gene expression play a crucial role in a wide variety of cellular processes. Dissecting the precise mechanisms of gene regulation is therefore necessary to understand both the normal functioning of cells and the ways in which dysregulation of certain genes plays a role in disease states^1^. Gene expression in metazoans is regulated by several distinct classes of *cis*-regulatory elements (promoters, enhancers, insulators, etc), with the activity of multiple enhancers being integrated to determine the expression levels of the average mammalian gene^2^. The activity of each enhancer or promoter element itself is driven by the concerted action of multiple DNA binding proteins called transcription factors (TFs), which typically bind to combinatorial grammars of short sequence motifs embedded in regulatory DNA sequences.

Functional genomic assays developed over the last decade (such as ChIP-seq, DNase/ATAC-seq, and others) have allowed for candidate *cis*-regulatory elements (cCREs) to be mapped on a genome-wide scale in a wide variety of cell lines and tissues^2,3^. They have more recently been supplemented by massively parallel quantitative measurements of the regulatory activity of native cCREs and synthetic constructs in the form of Massively Parallel Reporter Assays (MPRAs)^4–8^ and Self-Transcribing Active Regulatory Regions sequencing (STARR-seq)^9–13^ as well as direct high-throughput perturbations of cCREs in their native contexts using pooled CRISPR screens^14,15^.

However, determining the functional nucleotides and sequence patterns that drive regulatory activity in individual DNA elements remains challenging^16^. This is due to the difficulty in modeling how transcription factors bind to DNA, how their combinatorial binding activity is transformed into regulatory potential, and how multiple regulatory elements modulate transcriptional activity of target genes. Growing appreciation of the role of noncoding single nucleotide polymorphisms (SNPs) in numerous disease contexts^17^ makes resolving these challenges all the more urgent.

To address this problem, machine learning methods such as random forests, support vector machines (SVMs), and convolutional neural networks (CNNs) have been trained on functional genomics data to learn predictive models mapping DNA sequences to associated regulatory markers. Example outputs include transcription factor (TF) binding, gene expression, and alternative splicing^18–22^. Recently, these approaches have been used to model regulatory activity measurements from MPRAs. An SVM-based model^23^ was the top performer in a challenge that benchmarked several methods for predicting MPRA activity of DNA sequences flanking regulatory genetic variants^24^. Kalita *et al.*^25^ developed a statistical model to estimate allelic imbalance at regulatory variants based on MPRA measurements. Sample *et al.*^26^ and Bogard *et al.*^27^ recently introduced deep learning models of UTR sequences trained on MPRA datasets measuring polysome profiling and alternative polyadenylation respectively. Here, we present MPRA-DragoNN (Deep RegulAtory GenOmic Neural Network), a CNN-based framework to predict and interpret the transcriptional regulatory activity of noncoding DNA sequences as measured by MPRAs. We extend the work piloted by Paggi *et al.*^28^ to model the activity of 16,000 distinct regulatory regions in the K562 and HepG2 cell lines as measured by a specific MPRA design called Sharpr-MPRA^7^. We find that our model’s predictive performance is close to the moderate replicate concordance of the Sharpr-MPRA assay. We apply a feature attribution method called DeepLIFT^29^ to the trained CNN model, allowing us to infer each nucleotide’s contribution to the predicted MPRA activity of an arbitrary input sequence. This approach enables the identification of predictive TF motifs and grammars with cellular and genomic context-specific activity. Further, we evaluate the ability of the model to predict the allelic effects of single nucleotide polymorphisms (SNPs) in an independent MPRA experiment. We provide anecdotal examples showing how such predictions can supplement genome-wide association studies (GWAS) by isolating putative causal variants from larger lists of SNPs in linkage disequilibrium with one another. While focusing on Sharpr-MPRA data in this study, our approach is broadly applicable to other experimental designs, and we anticipate better prediction and interpretation accuracy when using more reproducible assays as input training data. Our study is a proof-of-concept that systematic interpretation of purported “black box” neural network models of regulatory DNA can offer a promising route towards improving our understanding of the regulatory code and the effects of noncoding genetic variation on molecular and disease phenotypes.

## 2. Methods

Our overall workflow consists of three major components. First, we train and optimize CNNs that predict regulatory activity of noncoding DNA sequences as measured by MPRAs. Next, we estimate the predictive contributions (importance) of individual nucleotides in input DNA sequences and compare these to DNA sequence features with known biological function. Finally, we present case studies focused on discovering novel regulatory sequence grammars and identifying putative functional genetic variants associated with gene expression variation. In this section, we first provide an overview of MPRA experiments, and then discuss the details of these steps.

### 2.1. MPRAs and the quantification of candidate enhancer activity

The objective of MPRA experiments is to quantify the regulatory activity of a large library of DNA sequences. This is typically accomplished by placing these sequences in DNA plasmids, a library of which is transfected into cells and regulatory activity is measured by high throughput sequencing of RNAs expressed from the plasmids. Most MPRA designs rely on placing synthetic constructs of 100-200 bp length upstream of a reporter gene, which in turn contains barcodes specific to each such construct. The parallel, high-throughput readout of activity of all input sequences is accomplished by sequencing these barcodes.

In this study, we specifically focus our analysis to the Sharpr-MPRA design introduced by Ernst *et al.*^7^ (**Figure 1A and B**). In the Sharpr-MPRA protocol, 15,720 295 bp-long regions centered on DNase-seq peaks in K562 and HepG2 human cells (an erythroleukemia and a hepatocarcinoma cell line, respectively) were tiled with a total of ~487K 145 bp-long fragments; each such 295 bp-long region is tiled by 31 sequences at 5 bp intervals. The library of 145 bp-long fragments was cloned upstream of either a minimal promoter (minP) or a strong (SV40P) promoter, with unique barcodes located within the reporter mRNA. Each of these two libraries was tested in both K562 and HepG2 cells, resulting in measurements across four different conditions in total (**Figure 1B**).

**Figure 1:**
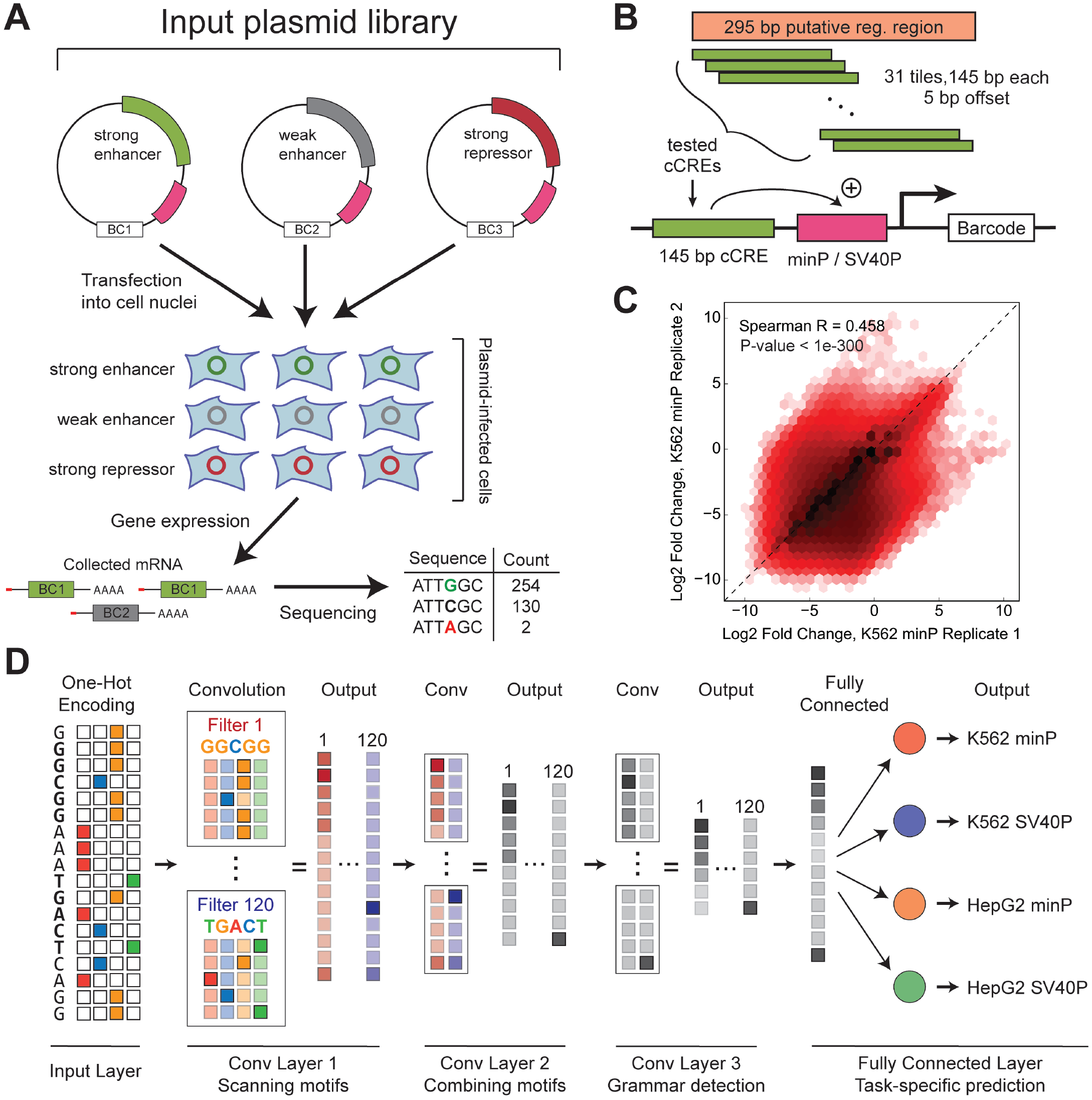
Predicting regulatory activity in MPRAs using convolutional neural networks. (**A**) Outline of the design of Sharpr-MPRA experiments used in this study. A collection of DNA constructs is cloned into a plasmid library upstream of a promoter (magenta) and transfected into a population of cells. Each construct is linked to a unique barcode (BC) located in the transcribed region; measuring the abundance of these barcodes using high-throughput sequencing allows for evaluation of the regulatory activity of each construct. (**B**) In the Sharpr-MPRA design, 145 bp-long 5-bp tilings of each of ~15,000 candidate 295 bp *cis*-regulatory elements are cloned upstream of either a minimal promoter (minP) or a strong promoter (SV40P). (**C**) Reproducibility between individual replicate Sharpr-MPRA measurements of regulatory activity (shown is data for K562 cells using the minP promoter). (**D**) Overview of the MPRA-DragoNN convolutional multi-task neural network architecture. The genomic DNA sequence for each tested MPRA construct is transformed from nucleotides (in ACGT alphabet) to a 145 × 4 one-hot encoded array. Three convolution layers and a fully-connected (FC) layer are then applied to predict four tasks (regulatory activity for the two cell lines with each of the two promoters). Each convolutional layer consists of 120 filters of length 5 (rectangles) that move along the sequence, searching for specific patterns of length 5 at every possible position. The first convolutional layer can be interpreted as identifying individual DNA sequence recognition motifs, such as those recognized by transcription factors. The second convolutional layer combines nearby potentially interacting motifs, while the third layer abstracts higher-order grammars (positioning, spacing, and other meta-features). Finally, the FC layer synthesizes these patterns with cell type– and promoter–specific information to make activity predictions.

While the between-replicate reproducibility of available Sharpr-MPRA data is modest (average Spearman correlation of enhancer activity across replicates = 0.40; **Figure 1C and S1**), this dataset has a key advantage in the fact that it is one of the larger MPRA studies. The large size of the dataset makes it a particularly good fit to train neural network models. Also, the uniform 5-bp tiling of cCREs allows for the contributions of individual nucleotides to regulatory activity to be evaluated more directly.

### 2.2. Training deep learning models to map DNA sequences to MPRA activity

We trained computational models to predict MPRA regulatory activity (training labels) from DNA sequence (training input) on the Sharpr-MPRA dataset. We chose convolutional neural networks (CNNs), a class of predictive models with state-of-the-art performance on tasks such as object recognition, language processing, and medical diagnosis^30^. CNNs are particularly effective at detecting spatial patterns in input data; as the salient features in regulatory sequences are thought to be specific combinations of consecutive base pairs (TATA boxes, TF motifs), CNNs are well-suited to the task of identifying key regulatory patterns.

We performed several standard data processing steps prior to training our models. Briefly, we (i) transformed the input data from length-145 ACGT strings to 145 × 4 “one-hot encoded” numerical arrays, in which an A corresponds to [1, 0, 0, 0], a C corresponds to [0, 1, 0, 0], etc; (ii) augmented our training dataset by adding the reverse complement of each original sequence, with the same output, as an additional example^31^; and (iii) *z*-score normalized regulatory activities within each task (mean 0, variance 1). Sequences from chromosomes 8 and 18 were held out for validation and testing respectively, with the remaining 457,174 examples (914,348 post-augmentation) composing the training set. Using separate whole chromosomes for training, validation, and testing ensures no overlap between sequences in the respective sets.

We experimented over a large search space to determine an optimal model architecture, varying the type, number, and numerical parameters of each CNN layer. We used the mean squared error of the predictions with respect to the experimental data for model optimization, a common choice for regression models. All models were trained in the Keras framework (version 1.2.2 with Theano backend) on an NVIDIA Tesla P100 GPU. Our final architecture (**Figure 1D**) consists of three convolutional layers (with ReLU activation) followed by a fully connected layer to predict the four tasks (K562 minP, K562 SV40P, HepG2 minP, HepG2 SV40P). We used a multi-task architecture in which a single model predicted all four tasks simultaneously, as this choice improved performance. Each convolutional layer has 120 scanning filters with length 5 followed by batch normalization and dropout (with *p*_dropout_ = 0.1), two well-established measures to reduce model overfitting.

Most of the hyperparameters we converged upon are generally consistent with recent literature in the field^32,33^, with two notable exceptions. First, we found that adding fully connected layers between the third convolutional layer and the final prediction layer reduced performance. Second, our optimal filter length was 5, which is smaller than previous CNN filter lengths used for genomics (usually 10-30). These findings suggest that model hyperparameters in functional genomics are application-specific and need to be optimized depending on the particular problem being studied.

### 2.3. Using DeepLIFT to estimate the predictive importance of individual nucleotides in regulatory DNA sequences

Deciphering the functional nucleotide patterns and grammars that are predictive of a DNA sequence’s regulatory activity is one of the main applications of our MPRA-DragoNN models. Given an input sequence, we want nucleotide-resolution importance scores quantifying each nucleotide’s contribution to the predicted output in a specific cell type. These scores then allow downstream analyses such as identifying transcription factor recognition motifs and combinations thereof, enabling formulation of specific biological hypotheses regarding the relationship between DNA sequence and regulatory activity.

Multiple methods have been developed to compute feature importance scores for CNNs. A common approach in genomics is *in silico* mutagenesis (ISM), in which the prediction for the reference input *f* (*X*_ref_) is compared to the prediction for mutated inputs *f* (*X*_mut_); the score for each nucleotide is the maximum difference *f* (*X*_ref_) − *f* (*X*_mut_) across the three possible mutations^21,33^. While intuitive, ISM is very computationally expensive, since it requires three forward passes through the network for each of the 145 nucleotides. Furthermore, noncoding sequences have been shown to have redundant features, *e.g.* two adjacent TF motifs with the presence of only a single one being necessary to drive gene expression; an accurate predictor would make equivalent predictions for the activity of the wild type sequence and all sequences with one of the two motifs mutated and thus ISM would fail to identify either of them as an important feature.

Hence, we used DeepLIFT, a recently developed backpropagation-based feature attribution method for neural networks that can estimate the predictive contribution (importance) of each nucleotide in an input sequence to its predicted output^29^. DeepLIFT requires a single backward pass through the network to compute contributions for all 145 nucleotides, making it orders of magnitude faster than ISM, and it has been demonstrated to overcome issues with ISM and similar methods^29^ of the kind described above. Throughout our analyses, we used dinucleotide-shuffled sequences (built-in implementation in the DeepLIFT package) as reference when computing DeepLIFT scores.

### 2.4. Other referenced datasets

In addition to Sharpr-MPRA, we referenced a number of other datasets to further understand and validate the performance of our model. We discuss the relevance of these data to MPRA-DragoNN in their respective Results subsections, but here we will briefly describe the datasets themselves.

To determine the chromatin state of the genomic regions from which the MPRA fragments were designed, we used annotations inferred by ChromHMM^34^. This method applies a Hidden Markov Model to learn a cell type-specific chromatin state for each 200 bp segment of the genome based on histone modification and chromatin accessibility datasets. We downloaded 25-state ChromHMM annotations for the K562 and HepG2 cell types generated by ENCODE^2^. We designated any fragments drawn from regions with “Tss” (active promoter) or “PromF” (promoter flanking) states as promoter fragments; “Enh”/“EnhF” (candidate strong enhancer) or “DnaseD”/“DnaseU”/“FaireW” (accessible weak enhancer) as enhancer fragments; and “Repr”/“ReprW”/“ReprD” (Polycomb repressed) or “Quies” (heterochromatin) as repressed fragments.

We referenced TF binding site (TFBS) predictions made using the CENTIPEDE algorithm^35^ in order to determine whether predictive features extracted by DeepLIFT agree with validated measures of biological function. Using histone modifications and DNase I footprints, CENTIPEDE fits a Bayesian mixture model to discriminate bound and unbound TF motif matches, generating fairly accurate genome-wide binding maps. We downloaded the set of K562 binding sites from www.centipede.uchicago.edu for our analysis.

To examine the generalization of our model’s predictions on external datasets not used during training, we chose the MPRA performed by Ulirsch et al.^36^. This experiment was designed to study the functional consequences of 2,756 GWAS variants associated with erythroid disorders, and was well-suited to our use for multiple reasons: (1) the assay was performed in K562 cells, allowing for comparison to our K562 prediction model; (2) the construct length used in it was 145 bp, the same length as in Sharpr-MPRA, facilitating the direct application of our model; (3) sequence activity measurements exhibited high reproducibility (between-replicate Pearson *r* = 0.85); and (4) most importantly, it tests the activity of both reference and mutant fragments to quantify the effects of sequence variants on regulatory activity. Thus, taking these data as gold standard, we evaluated our model’s ability to predict the effect size of the expression change induced by such variants (described in the Results).

We also tested our model’s ability to prioritize and fine-map variants identified by genome-wide association studies (GWAS). For this evaluation, we used the dataset from Willer *et al.*^37^, in which the genotypes of 2,437,752 SNPs were correlated with fasting blood lipid levels in 189,000 European and 8,000 non-European individuals. We specifically focused our analysis to the strength of association with low-density lipoprotein (LDL) levels, a well-studied risk factor of cardiovascular disease and myocardial infarction^38^.

## 3. Results and Discussion

### 3.1. MPRA-DragoNN predicts measured MPRA regulatory activities on par with replicate concordance of the assay

We evaluated the performance of our models by computing the Spearman correlation between experimentally measured regulatory activities (averaged across replicates) and our model’s regulatory activity predictions. Because Spearman correlation only depends on the ranks of the data, it is less susceptible to artificial performance inflation due to outliers (as is Pearson’s *r*) or center-heavy distributions (mean squared error).

The Spearman correlation between predicted and experimentally measured values for the held-out testing set was 0.28 **(Figure 2A)**(and 0.14, 0.21, and 0.22 for K562 SV40P, HepG2 minP, and HepG2 SV40P, respectively; **Figure S2A-C**). These values are low in absolute terms, but in the context of the relatively weak replicate concordance of the assay itself (**Figure 1C and S1**), the performance suggests that the model captures much of the putative biological signal present in the data. For K562 minP, 61% (the ratio between the respective Spearman correlations) of experimental reproducibility is accounted for by our model predictions (averaged for the four tasks, this ratio is 52%). We also found that the model’s prediction error for a given DNA sequence was positively correlated with the difference in its two replicate activity values (Spearman *ρ* = 0.29, *P* < 10^−181^, **Figure S2D**), suggesting that more reproducible data is met with more accurate prediction.

**Figure 2:**
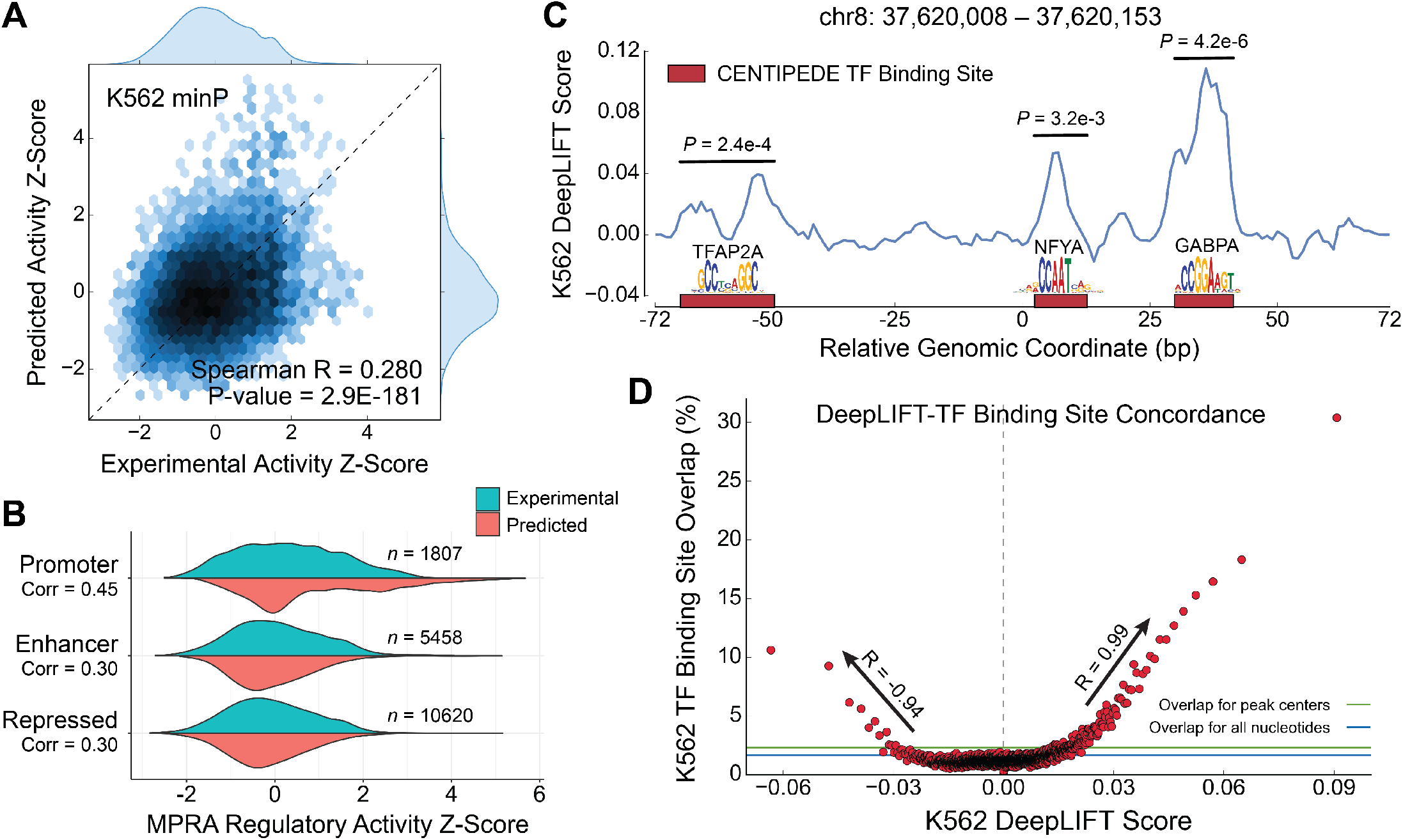
MPRA-DragoNN distinguishes active regulatory sequences at high resolution. (**A**) Predicted regulatory activity *z*-scores vs. experimental activity *z*-scores for the K562 minP task. (**B**) Distributions of experimental and predicted regulatory activities for different ChromHMM-inferred chromatin states. (**C**) K562 DeepLIFT nucleotide score track for a strongly activating regulatory sequence (top 0.1%) containing three TF binding sites (red) as identified by the CENTIPEDE algorithm. All three TFBSs are detected with statistical significance (Mann-Whitney *U* test). (**D**) Nucleotides with strong (in absolute value) DeepLIFT scores are more likely to overlap with TF binding sites than control sequences (blue: all nucleotides, green: DNase peak centers). This trend holds for both positive (*R* = 0.99) and negative scores (*R* = −0.94).

We then examined prediction performances (for the K562 minP task) on subsets of the testing set, with each subset consisting of all the 145 bp fragments from either promoter, enhancer, or repressed chromatin states. As shown in **Figure 2B**, model predictions generally appear to follow similar distributions as the experimental values for each of these three states. We observe markedly higher performance (Spearman *ρ* = 0.45) for fragments within or flanking gene promoters, suggesting that these sequences either are more experimentally reproducible or have more consistent predictive patterns that the model can learn. Constructs in DNase- or FAIRE-accessible regions were also particularly well-predicted, with an experiment-prediction Spearman *ρ* = 0.57 (**Figure S2E**).

### 3.2. Analysis of predictive nucleotides inferred from the model

Applying DeepLIFT to our top-performing MPRA-DragoNN model, we computed contribution scores for 4 million nucleotides lying in K562, HepG2, MCF-7, or HeLa DNase-seq peaks (27,886 total peaks from held-out chromosomes 8 and 18; each peak was clipped to 145 bp). We examined concordance between these scores and validated markers of regulatory function, such as putative TF binding sites (TFBS). Anecdotally, most of the strongly predicted sequences had high DeepLIFT importance scores at CENTIPEDE-annotated TFBSs. **Figure 2C** shows an example DeepLIFT score profile for a particular 145 bp fragment containing three CENTIPEDE-defined TFBSs; each of the sites is identified at fine resolution, with nucleotides within the binding sites assigned higher importance scores than the rest of the bases in the region (*p* < 5 × 10^−3^, Mann-Whitney *U* test).

To evaluate concordance between DeepLIFT scores and TFBSs on a broader scale, we sorted nucleotides by their DeepLIFT scores in K562 and binned them into 1,910 quantiles of 2,117 bases each. We find that DeepLIFT scores correlate strongly with TFBS overlap in K562, with > 30% of nucleotides in the highest DeepLIFT quantile overlapping a CENTIPEDE-annotated TFBS **(Figure 2D)**. Notably, this score-overlap relationship holds both for positive DeepLIFT scores (*R* = 0.99; *P* = 3.3 × 10^−72^, for scores higher than +0.2) and negative scores (*R* = −0.94; *P* = 2.2 × 10^−29^ for scores lower than −0.2), suggesting that of our model has the potential to identify both activating and BRCA1, and CHD2 in multiple databases, however, repressive motifs that modulate gene expression in concert.

### 3.3. Predictive sequence patterns are enriched at motifs of lineage-specific transcription factors

We performed several further evaluations of the ability of our model’s DeepLIFT scores to identify putative regulatory sequence patterns. We first compared DeepLIFT scores with nucleotide-level regulatory scores as calculated in the Sharpr-MPRA study itself, expecting considerable overlap as both methods use the same data. To do so, we referenced a compendium of mapped known TF binding motifs from Kheradpour *et al.*^39^. We separately averaged the DeepLIFT and Sharpr scores (from K562 cells) for the 4-20 base pairs comprising each motif match (a total of ~328,083 matches spanning 1,934 TF motifs), and evaluated whether the mean was significantly higher than that of negative control shuffled matches using a *z*-test.

Out of the ~328,000 motif matches, 51,730 were deemed significant by at least one of the methods at a Benjamini-Hochberg corrected false discovery rate (FDR) threshold of 0.1. Notably, 10,556 (20.4%) of these matches were identified by both DeepLIFT and Sharpr (**Figure 3A**), a 4.03-fold enrichment over the number of overlapping motifs expected by chance if the two methods were independent. Across all ~328,000 motif instances, Sharpr and DeepLIFT scores also agreed with Pearson *r* = 0.43 (**Figure S3A**).

**Figure 3:**
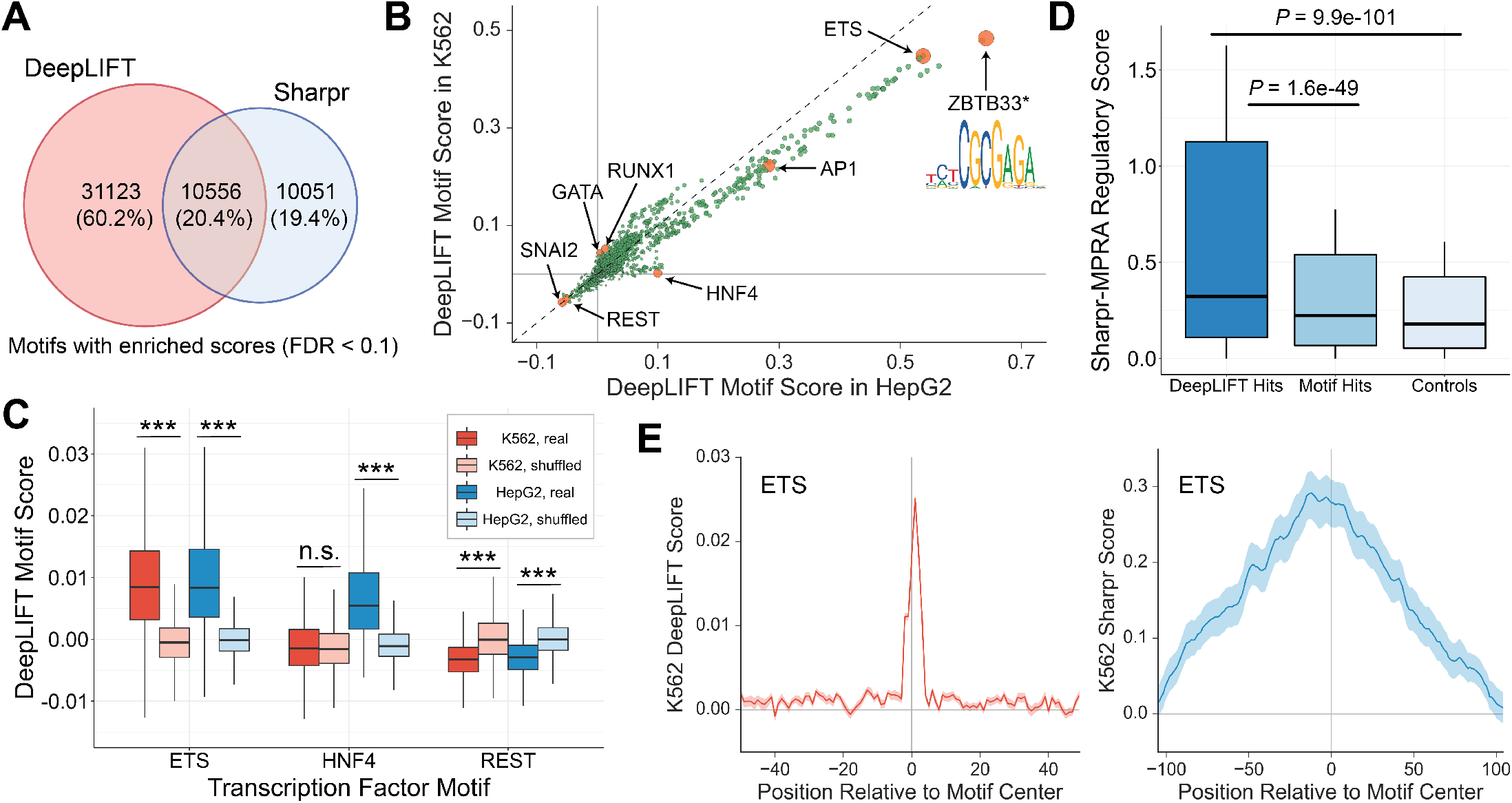
MPRA-DragoNN DeepLIFT feature importance scores robustly predict functional nucleotides. (**A**) Overlap between significant motif instances (Benjamini-Hochberg FDR < 0.1) identified by DeepLIFT and Sharpr. (**B**) Scatter plot of average DeepLIFT scores for 1934 motifs in HepG2 (*x*-axis) and K562 (*y*-axis)^39^. Orange points are discussed in the text. (**C**) Sharpr-MPRA nucleotide score distributions for (i) motifs that are also DeepLIFT hits, (ii) all motifs, and (iii) negative control shuffled motifs. (**D**) Distributions of average DeepLIFT motif scores for ETS, HNF4, REST, and their respective control motifs (shuffled versions) in both K562 and HepG2. \*\*\**p* < 10^−200^; n.s., not significant. (**E**) Positional distribution of DeepLIFT scores (left) and Sharpr scores (right) with respect to the center of ETS motif occurrences. Note that the DeepLIFT plot *x*-axis ranges from −50 bp to 50 bp while the Sharpr plot ranges from −100 bp to 100 bp. All *p*-values are computed with the Mann-Whitney *U* test.

We then computed consensus scores for each of the 1,934 TF motifs by averaging scores across all of the motif’s matches. Because these scores are each derived from on average ~170 motif matches, they tend to be less noisy and are thus more strongly correlated with their counterpart Sharpr scores (Pearson *r* = 0.86, **Figure S3B**). We examined the top scoring motifs from this analysis, and whether they included transcription factors known to be important in each of the two cell lines (**Figure 3B**). Consistent with known roles, ETS and AP-1 were identified as strong activators while the well-studied transcriptional repressors REST and SNAI2 had among the most negative scores in both K562 and HepG2 cells^40^. DeepLIFT also accurately retrieved the cell-type specificities of TFs: HNF4, a critical TF for liver development^41^, scores highly in HepG2 but not K562, while GATA1/2 and RUNX1, TFs with known roles in blood cells^42^, are specific to K562 cells. We further confirmed that these motifs indeed represent biologically relevant findings instead of noise by comparing the distributions of scores across all motif instances of ETS, HNF4, and REST to their respective control (instances of shuffled versions of the motifs) distributions (**Figure 3C**).

Interestingly, the top scoring motif in both cell types, a palindromic TCTCGCGAGA pattern (**Figure 3B**), is not a well-characterized protein binding motif. *In vitro* experiments and limited *in vivo* evidence suggest association with the zinc finger protein ZBTB33^43^; we also note that the motif is listed as associated with proteins such as DYRK1A,^39,44^, however, this is most likely a case of spurious annotations as these are not sequence-specific DNA binding proteins. Regardless of which protein(s) it is bound by, its high predictive value according to DeepLIFT suggests that instances (and disruptions) of this sequence may have substantial effects on expression.

We also found that motifs with statistically enriched DeepLIFT profiles (FDR < 0.05 when comparing the motif’s DeepLIFT scores to the shuffled motif DeepLIFT score distribution) have significantly higher Sharpr activity than the unfiltered set of all motifs (*P* = 1.6 × 10^−49^, Mann Whitney *U* test), supporting our model’s ability to identify active nucleotides (**Figure 3D**).

Another important consideration is the resolution at which the model identifies functionally relevant nucleotides. To address that question, we used all ~4000 instances of the strongly enhancing ETS motif, and computed the mean DeepLIFT score for each position relative to the motif center. As shown in **Figure 3E**(left), DeepLIFT perfectly highlights the core 6 bp CCGGAA motif, with virtually no signal for the surrounding base pairs. In contrast, the Sharpr score track contains a peak near the motif center, but the scores are still enriched for the surrounding 200 bp region, highlighting the finer resolution of MPRA-DragoNN.

Taken together, our results demonstrate that, genome-wide, our model’s DeepLIFT scores tend to prioritize putative regulatory nucleotides.

### 3.4. Inferring active transcription factors from nucleotide scores

Having confirmed that our model’s DeepLIFT feature importance scores can broadly identify functionally relevant TFBSs, we sought to apply them to understand the regulatory properties of the sequences tested in the Sharpr-MPRA. Using our nucleotide-resolution regulatory activity contributions, we aimed to determine which transcription factors confer enhancer function to each 145 bp DNA sequence. To do so, we downloaded position-weight matrices (PWMs) for 344 TF motifs from the HOMER database^45^. For each of the ~974,000 sequences in the Sharpr dataset, we assigned a score for how well each TF’s PWM aligned with the sequence’s DeepLIFT track (i.e., the maximum dot product between the PWM and the DeepLIFT track across all possible positions in the sequence; this metric accounts for both the magnitudes of the DeepLIFT scores as well as for how closely the PWM matches the sequence, for a combined causality estimate of how much a TF’s PWM contributed to the CNN’s prediction). It’s important to note that these scores are largely driven by underlying motif occupancy in a sequence; the DeepLIFT profiles act as a ‘multiplier’ to quantify how much a motif actually affects activity. This computation results in a ~974, 000 × 344 matrix, where element (*i, j*) can be interpreted as motif *j*’s “usage score” in controlling the expression of sequence *i*. We averaged across the two cell types.

First, we considered whether promoter and enhancer regions are distinguished by the TFs that they use to modulate gene expression. For each TF, we computed its average usage scores across all annotated promoters and across all annotated enhancers, and calculated the ratio of its promoter usage to its enhancer usage (**Figure 4A**). We found that motifs for TFs in the Sp/KLF C2H2 zinc finger sub-family are most specific to promoter regions, consistent with previous results^46,47^; other motifs enriched in promoters include NRF1, ZBTB33, and c-Myc, as well as ETS-related motifs. All of the most enhancer-specific TFs are in the AP-1-like subfamily of bZIP transcription factors. Interestingly, however, CREB, a bZIP factor, is a strongly promoter-enriched TF, in agreement with previous work highlighting its functional differentiation from AP-1^48^.

**Figure 4:**
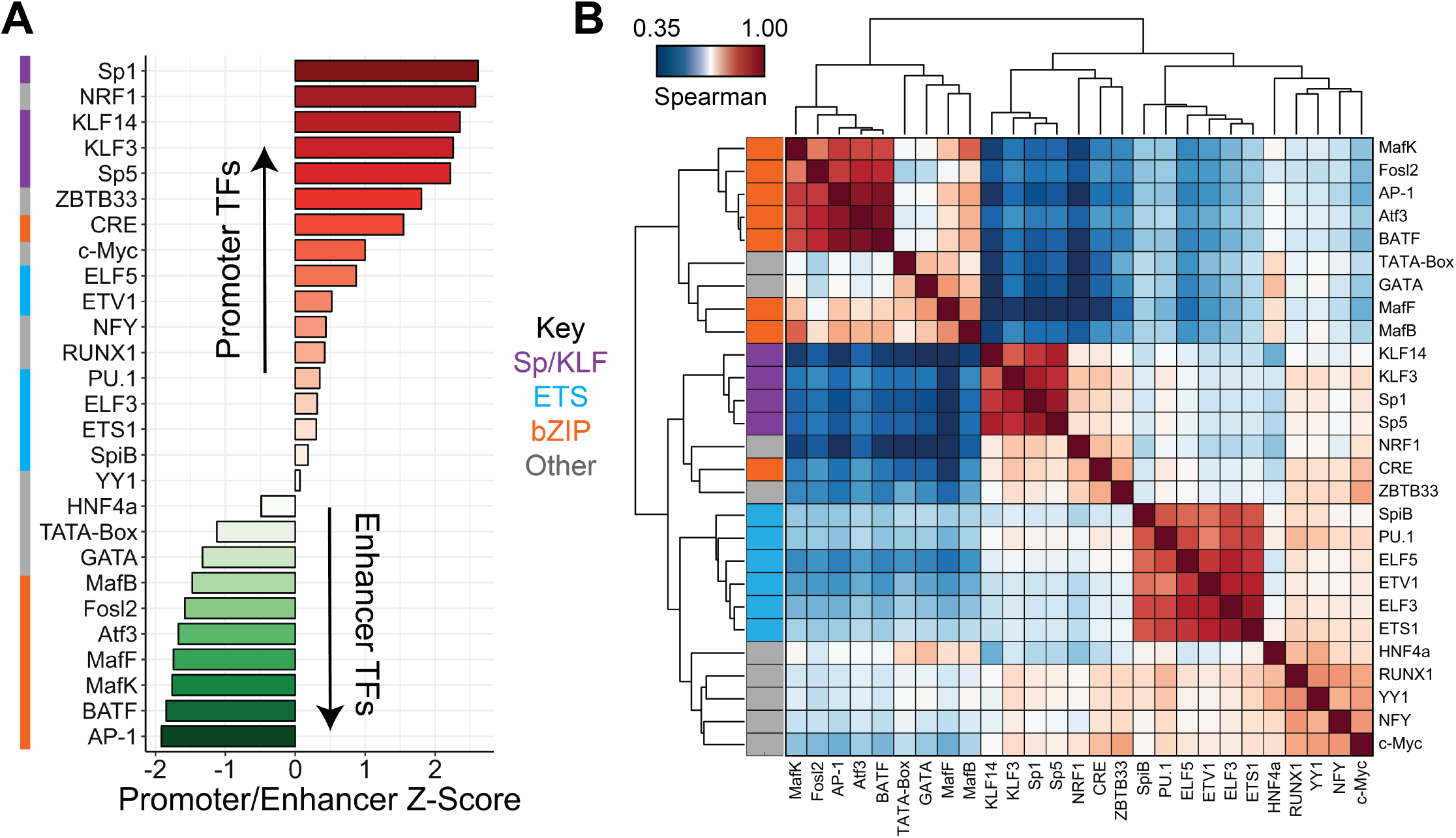
MPRA-DragoNN reveals patterns of transcription factor activity. (**A**) For each TF, we computed the ratio of average “usage” in promoter sequences relative to enhancer sequences. The plot contains *z*-scores of this ratio for 27 selected transcription factors, colored by their motif family (left). (**B**) Clustered correlation matrix of TF usage for the 27 factors from (A). Each cell is colored according to the motif usage Spearman correlation for a given pair of TFs across all ~974,000 sequences. Rows are colored by their motif family.

We next characterized correlated pairs of predictive TF motifs across the entire dataset (**Figure 4B**). As expected, we find that TFs within the same family cluster together; however, many of these correlations might be accounted for by the similarity of the underlying motifs rather than true co-occurrence of distinct motifs. The remaining TFs separate into two clusters, largely differentiated by promoter or enhancer specificity as discussed above.

More interesting are observed correlations between TFs of different families. For example, we find that HNF4 and GATA motifs are strongly correlated; indeed, previous work has found that the two TFs can have coordinated functions^49^. Another example involves MafB and MafK, which, despite different binding motifs, also correlate well, supporting previously predicted heterodimerization between the two^50^. In contrast, some TF pairs are very weakly correlated. For example, Sp/KLF factors do not correlate with the bZIP family (AP-1-like), possibly reflecting their differential promoter-enhancer preferences **Figure 4A**.

### 3.5. MPRA-DragoNN predicts allelic effect of genetic variants tested in an independent MPRA experiment

We next aimed at establishing to what extent our model’s predictions generalize beyond the Sharpr-MPRA dataset. To this end, we used MPRA data from Ulirsch et al.^36^, in which the regulatory activity of a number of reference sequences and mutated versions of each of these sequences were experimentally tested using MPRAs in K562. By computing the change in activity between the mutated and reference sequences, the authors quantified the regulatory impact of 8K variants, of which 40 had statistically significant effects. We generated analogous predictions for each of these 40 reference and mutant sequences, calculated the differences in predictions, and then compared our model’s inferred variant effects to the actual experimental data. We chose the *in silico* mutagenesis (ISM) approach over DeepLIFT in this case, as we were specifically interested in how alleles of individual SNPs affect predicted reporter expression.

We find that variant ISM scores are well-correlated with experimentally measured variant scores (**Figure 5A**). Of the 40 variants examined, our model correctly identifies the direction of effect (*i.e.*, increase or decrease in expression) for 32 of them. We explored one of the SNPs, rs2269907, that our model correctly predicted to increase expression. As shown in **Figure 5B**, this variant is located in an active candidate enhancer region and also lies within a ChIP-seq peak for the JunD transcription factor^51^. The variant appears to be correcting a mismatch in the JunD motif, likely causing JunD to bind and positively regulate gene expression.

**Figure 5:**
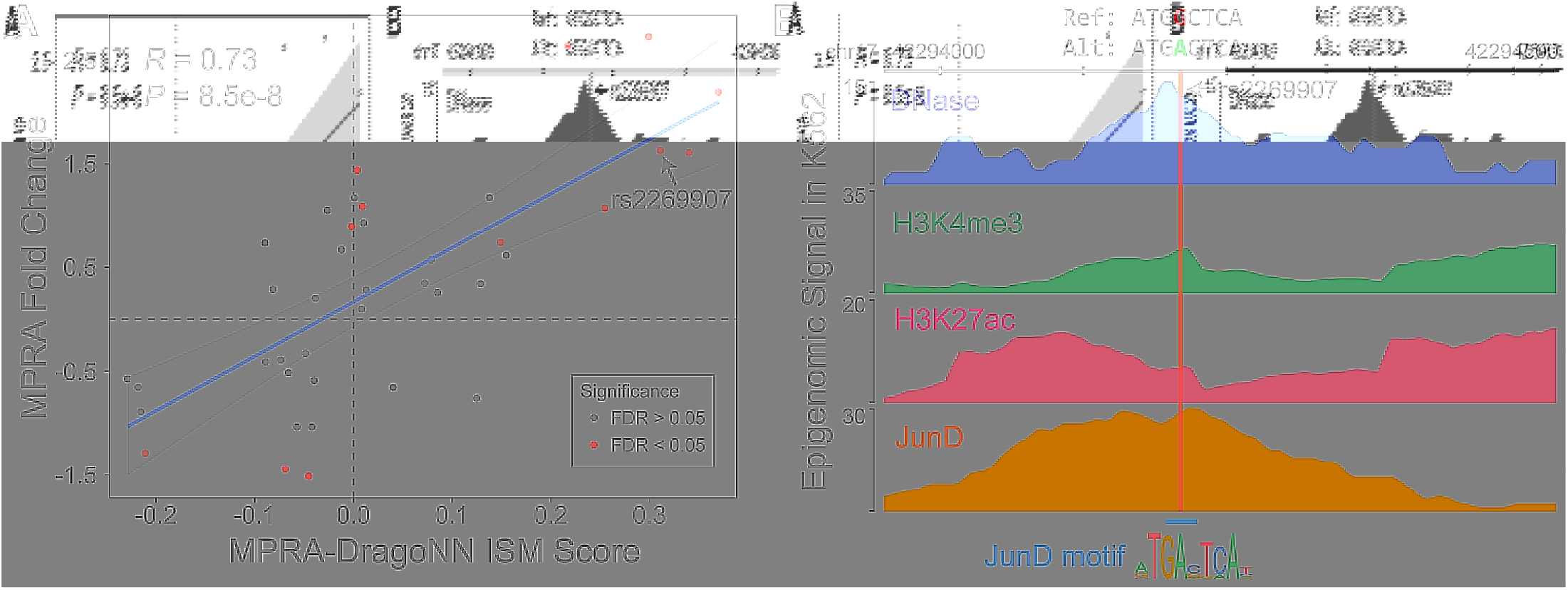
Variant in-silico mutagenesis scores agree with experimental data. (**A**) Regulatory activity changes between reference and mutated sequences predicted by MPRA-DragoNN agree with experimentally measured changes^36^. Red points indicate variants that were significant in the wild-type K562 condition (see description in Methods) (**B**) Detailed examination of a particular variant, rs2269907 at chromosome 17 position 44,294,214. The distribution of epigenetic marks and JunD ChIP-seq signal^2^ around the variant reveals that it lies in an active region^51^; the variant appears to correct a mismatch in one of the base pairs of a JunD motif, allowing JunD to bind and regulate expression.

These observations suggest that our model learns regulatory patterns that can generalize to contexts other than the specific MPRA dataset it was trained on.

### 3.6. MPRA-DragoNN can prioritize genetic variants associated with lipid traits

Genome-wide association studies (GWAS) have identified thousands of noncoding genetic loci associated with hundreds of complex traits and diseases. Noncoding disease-associated genetic variants tend to be enriched in regulatory elements and affect regulatory molecular phenotypes such as TF binding, chromatin accessibility and gene expression^2,52,53^. However, identifying the causal variants within disease-associated loci is often challenging due to linkage disequilibrium between multiple highly correlated SNPs in each locus^54^. We wanted to test whether MPRA-DragoNN could prioritize likely causal variants within complex trait-associated loci based on their predicted impacts on MPRA regulatory activity.

Complex diseases that are influenced by gene misregulation are often cell-type specific. We therefore chose traits associated with liver, since we could leverage our model’s predictions in the HepG2 liver carcinoma cell type. We referenced a GWAS in which genotypes at ~2.4M loci were analyzed to find SNPs significantly correlated with LDL cholesterol levels^37^. We then scored these SNPs using the *in silico* mutagenesis (ISM) procedure described in Section 2.3. That is, we generated a prediction for the reference sequence, containing the wild type allele of the SNP and 72 base pairs of surrounding context upstream and down-stream (1 + 2 · 72 = 145, length of the inputs to our model), and we also generated mutated predictions for the three possible nucleotide changes at the location of the SNP. Each nucleotide’s ISM score is the greatest absolute value difference from the reference prediction across the three mutants (as we did not always know the specific ref/alt alleles of the SNPs in our dataset). We again used ISM instead of DeepLIFT as we were specifically interested in testing changes to predicted expression as a result of individual sequence variants.

We expanded the dataset of GWAS-tested SNPs (tag SNPs) to include all other SNPs in LD (*R*^2^ ≥ 0.8) with them. We scored both the tag SNPs and all of their proxy SNPs, and each tag SNP was ultimately assigned an LD-adjusted score: *i.e.*, the maximum ISM score over all the variants in its LD block.

Overall, we find that statistically significant tag SNPs with GWAS *p*-values less than 5 × 10^−8^ have higher LD-adjusted ISM scores than the LD-adjusted ISM scores of insignificant (*p* > 0.1) variants (Mann-Whitney *U* test: *p* = 8.8 × 10^−9^; **Figure S4A**). The effect size of this difference, however, is relatively small, with the former set having a 15% higher mean score than the latter set. To tell whether the LD score adjustment improved our discriminative power, we compared unadjusted ISM scores for significant tag SNPs to the unadjusted scores of insignificant tag SNPs. There was no difference between these two distributions of scores (*p* = 0.60; **Figure S4B**), suggesting that the tag SNPs tested in GWAS usually aren’t driving their predicted regulatory effects; their proxy SNPs must be considered to elucidate their putative function. We also observe a low but significant negative correlation between LD-adjusted ISM scores and *p*-values of the corresponding tag SNPs (*ρ* = −0.07, *p* = 3.1 × 10^−37^); that is, higher ISM scores are weakly associated with lower GWAS *p*-values (**Figure S4C)**. It is worth noting that predicting trait-association of variants from a model trained to predict MPRA activity or any regulatory molecular marker is a very challenging task. That is, variants with larger effects on expression do not necessarily have lower GWAS *P*-values than other variants, and vice-versa. Models trained on higher quality MPRA datasets and adaptively fine-tuned on trait-association statistics may exhibit improved performance on this prediction task.

We next wanted to test whether combining information from GWAS with MPRA-DragoNN predictions could better prioritize variants. We plotted each SNP with its ISM score on the *x*-axis and its negative log GWAS *p*-value on the *y*-axis (**Figure 6A**). In this setup, putatively causal variants localize to the upper right or upper left quadrants, as those points have both large effects on predicted expression and strong correlation with disease phenotypes. We focused on SNPs with absolute value ISM scores above 0.45 and *p*-values below 5 × 10^−8^, which revealed 263 variants associated with low-density lipoprotein levels. Most of the top mutations occurred proximally upstream or within introns of previously implicated cholesterol/cardiovascular disease genes, including *LPIN3*, *FADS1/2*, *HLA-C* and *APOB*^38,55–57^.

**Figure 6:**
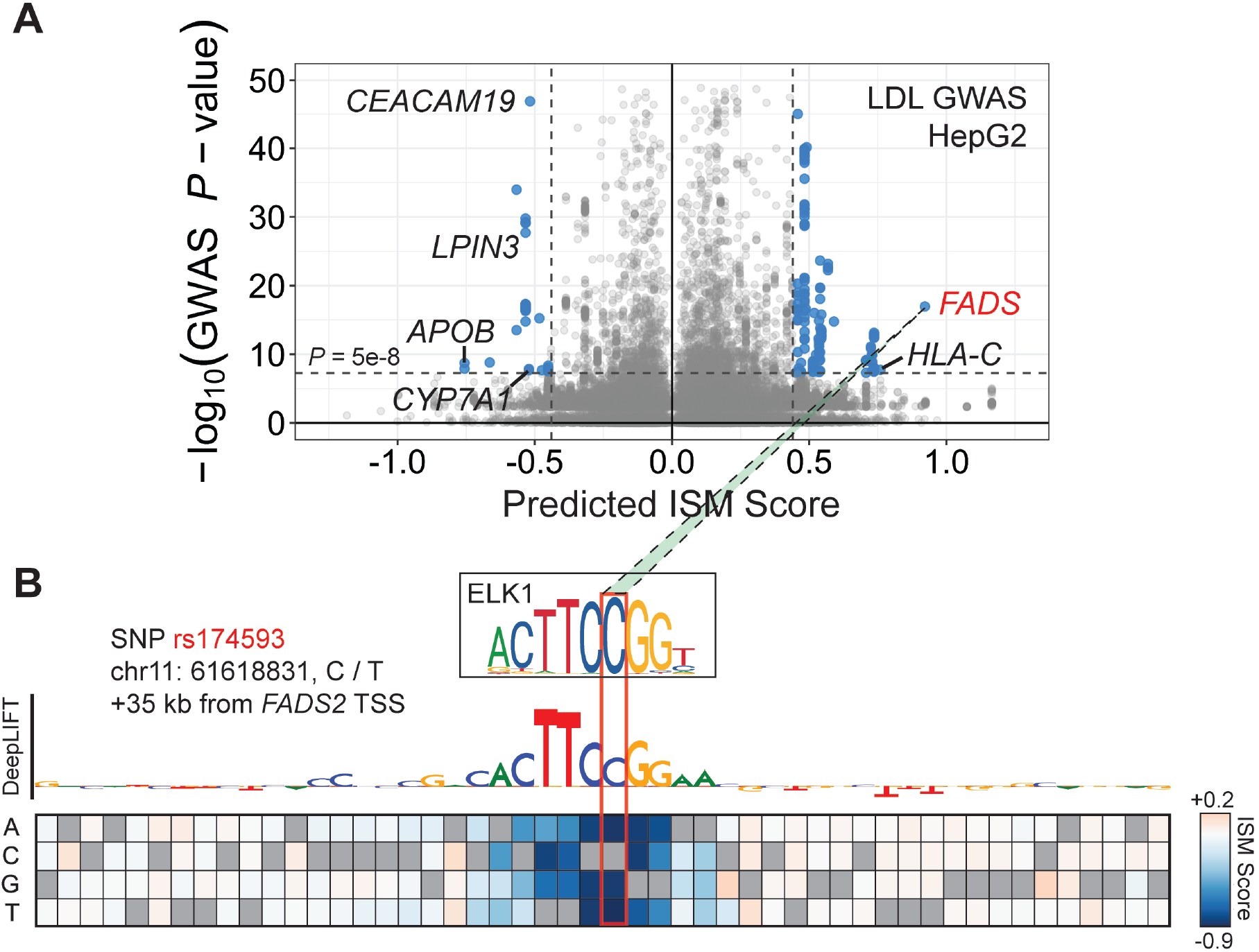
Dissecting rs174593, a putative causal variant for reduced LDL cholesterol levels. (**A**) Volcano plot with in-silico mutagenesis scores on the *x*-axis and negative log GWAS *p*-values on the *y*-axis. Putatively causal variants in the upper right and upper left regions localize to cardiovascular disease-related genes. (**B**) DeepLIFT track and saturation mutagenesis scores of the locus surrounding rs174593, a potential *FADS2 cis*-regulatory element. As highlighted by DeepLIFT, the C allele at that position creates an ELK1 motif match, increasing predicted *FADS2* expression compared to the T allele.

We focused on the variant rs174593 (in LD with tag SNP rs174591) at the *FADS* locus, which is predicted to result in a +0.92 change in regulatory activity *z*-score and has a GWAS *p*-value of 10^−17^. This SNP lies in an intron of *FADS2*, which codes for fatty acid desaturase 2, an enzyme that converts long-chain saturated fatty acids to polyunsaturated fatty acids (PUFAs)^58^. Notably, PUFA levels are inversely correlated with blood LDL content^59^. To further investigate the putative mechanisms through which rs174593 influences gene expression, we computed DeepLIFT scores of surrounding nucleotides and also performed a complete saturation mutagenesis of the 25 base pairs on either side of the SNP (**Figure 6B**). As shown in **Figure 6B**, the mutant C allele of the variant creates a near-perfect match to the strongly activating ELK1 transcription factor motif **(Figure 3B)**, potentially causing ELK1 to bind and increasing *FADS2* expression. While **Figure S5A** suggests only modest chromatin activity at the rs174593 locus in HepG2 cells^3^, tissue expression data from GTEx^60^ implicate rs174593 as an eQTL in liver cells with 92% posterior probability (**Figure S5B-C**). Further, ChromHMM annotations using Roadmap Epigenome^3^ data predict rs174593 to be located in the “Enhancer” or “Genic enhancer” states for multiple tissues (Heart, Testis, Esophagus, among others) in which *FADS2* is expressed and rs174593 is a *FADS2* eQTL (GTEx; **Figure S5C**), suggesting that the variant may influence disease through its effects on non-liver tissues as well. As higher *FADS2* expression increases PUFA production and therefore decreases plasma LDL levels, the mutant C allele of rs174593 may have a protective effect by reducing risk of LDL cholesterol-mediated atherosclerosis^61^.

While rs174593 is only an anecodotal example, the fine-mapping approach described here may become systematically successful when integrated with higher-quality datasets and more accurate models of multiple regulatory phenotypes. Our preliminary work hints at how interpreting predictive models can identify putative mechanisms through which disease mutations exercise their influence rather than simply identifying SNPs that are only correlated with disease, therefore multiplying the utility of thousands of existing genome-wide association studies.

## 4. Discussion

In recent years, functional genomic assays such as the numerous methods for profiling chromatin features, MPRAs, and pooled CRISPR perturbation screens have produced genomic data at unprecedented breadth, depth, and detail. MPRAs in particular present a highly scalable platform for finely dissecting the regulatory code of individual noncoding DNA elements, as they allow for large numbers of short sequences to be tested in parallel and in diverse cellular contexts. The expression-based readout of MPRAs is complementary to other assays like ChIP-seq (protein binding) and DNase-seq (chromatin accessibility), which do not directly measure effects on gene expression. Predictive models trained on MPRAs are hence more likely to be sensitive to identifying functional regulatory patterns that affect gene expression.

The increasing size and design complexity of MPRAs in the literature motivated us to develop MPRA-DragoNN, a CNN-based predictive model for learning *de novo* regulatory patterns from noncoding DNA sequences based on their MPRA activity. We applied our model to the Sharpr-MPRA dataset, demonstrating that we can predict quantitative regulatory activity with moderate accuracy within the range of replicate concordance of the assay. We then used state-of-the-art model interpretation approaches such as DeepLIFT and ISM to decipher the predictive sequence features learned by the model which correspond to motifs of contextually relevant transcription factor complexes. We demonstrated our model’s ability to generalize and predict the impact of genetic variation on regulatory activity measured in independent MPRA experiments. Finally, we tested MPRA-DragoNN’s ability to predict variants associated with complex traits from GWAS studies. Using LDL cholestrol GWAS as a case study, we found that variant ISM scores were only weakly correlated with association statistics even after accounting for LD. However, by combining GWAS summary statistics with model predictions, we were able to prioritize some candidate causal variants. Our case study of SNP rs174593 showcased how interpretation methods such as DeepLIFT and ISM can provide hypotheses about regulatory mechanisms of a putative causal variant. By integrating other widely available epigenomic datasets (DNase-seq, histone ChIP-seq) with higher-quality MPRA data, we expect to train far more accurate gene expression predictors; used in the way we preview here, these models may play a significant role in the quest towards mapping the relationship between sequence variants and gene expression changes in the context of human disease.

We conclude with a discussion of limitations of our study as well as general caveats associated with interpreting regulatory models trained on MPRAs. In this study, our primary goal was to explore the application of neural network models trained on MPRA data to interpret regulatory DNA sequences and noncoding variation. We have not included a systematic comparison of our CNN-based model to strong baseline predictors such as SVMs. We plan to focus on rigorous model comparisons in the future. There are also several feature attribution methods for interpreting neural network models. Here, we have used two state-of-the-art methods namely DeepLIFT and ISM. In the future, we hope to perform systematic comparisons of other interpretation methods^18^; study the stability and uncertainty of discovered patterns across multiple bootstrapped models and model architectures; and use improved methods for summarizing globally predictive patterns^26,27^ and higher-order feature interactions^62^. Further, the moderate replicate concordance of the Sharpr-MPRA assay presented a significant challenge to train a high-fidelity model. While MPRAs can test hundreds of thousands of sequences simultaneously, we have found several published datasets to exhibit high variance in replicate measurements (results not shown), limiting the ability to train reliable models. We expect that data quality will improve with further optimization of the assays resulting in more accurate models. In addition, MPRAs typically test relatively short sequences that likely do not encompass complete regulatory elements. Finally, most MPRAs have been performed using episomal DNA plasmids rather than native chromatin, leaving open the question to what extent their output is directly translatable to the native genome, in which overall chromatin context plays a significant role in determining regulatory output. Using models trained on MPRAs alone as variant effect predictors may be ill-advised, as we observed many SNPs disrupting regulatory sequences that are active in MPRAs but endogenously repressed (and vice-versa). Some of these limitations are being addressed, as long-fragment (~500 bp) STARR-seq datasets^10^ and genome-integrated MPRAs^63^ have become recently available. MPRA-DragoNN can easily adapt to these assays. Transfer learning approaches may also be able to boost predictive models of *in vivo* transcription factor binding, chromatin state, and gene expression by pretraining on MPRA experiments. In conclusion, neural network models trained on a diverse collection of high-quality datasets coupled with powerful interpretation frameworks have the potential to finely decode *cis*-regulatory grammars and functional genetic variation in regulatory DNA sequences.

## Availability

All the code to reproduce the analyses described in the paper is available at https://github.com/kundajelab/MPRA-DragoNN/. The model is also uploaded to Kipoi (https://kipoi.org/), allowing users to generate activity predictions and feature importance maps for arbitrary 145-bp sequences.

## Author Contributions

RM and AK conceived of the project. RM and SN developed software. RM performed all analyses with advice from PG, GKM, and AS. RM drafted manuscript, and AK, PG, and GKM edited the manuscript.

## Acknowledgments

We thank several members of the Kundaje lab for their feed-back and suggestions on improving prediction performance and validating the model.

## Funding

PG was supported by a Stanford Interdisciplinary Graduate Fellowship. GKM was supported by the Stanford School of Medicine Dean’s Postdoctoral Fellowship. AS was supported by an HHMI International Student Research Fellowship and a Bio-X Bowes Fellowship. AK was supported by National Institute of Health grants 1DP2GM123485 and 1U01HG009431.

## Supplementary Figures

**Supplementary Figure 1:**
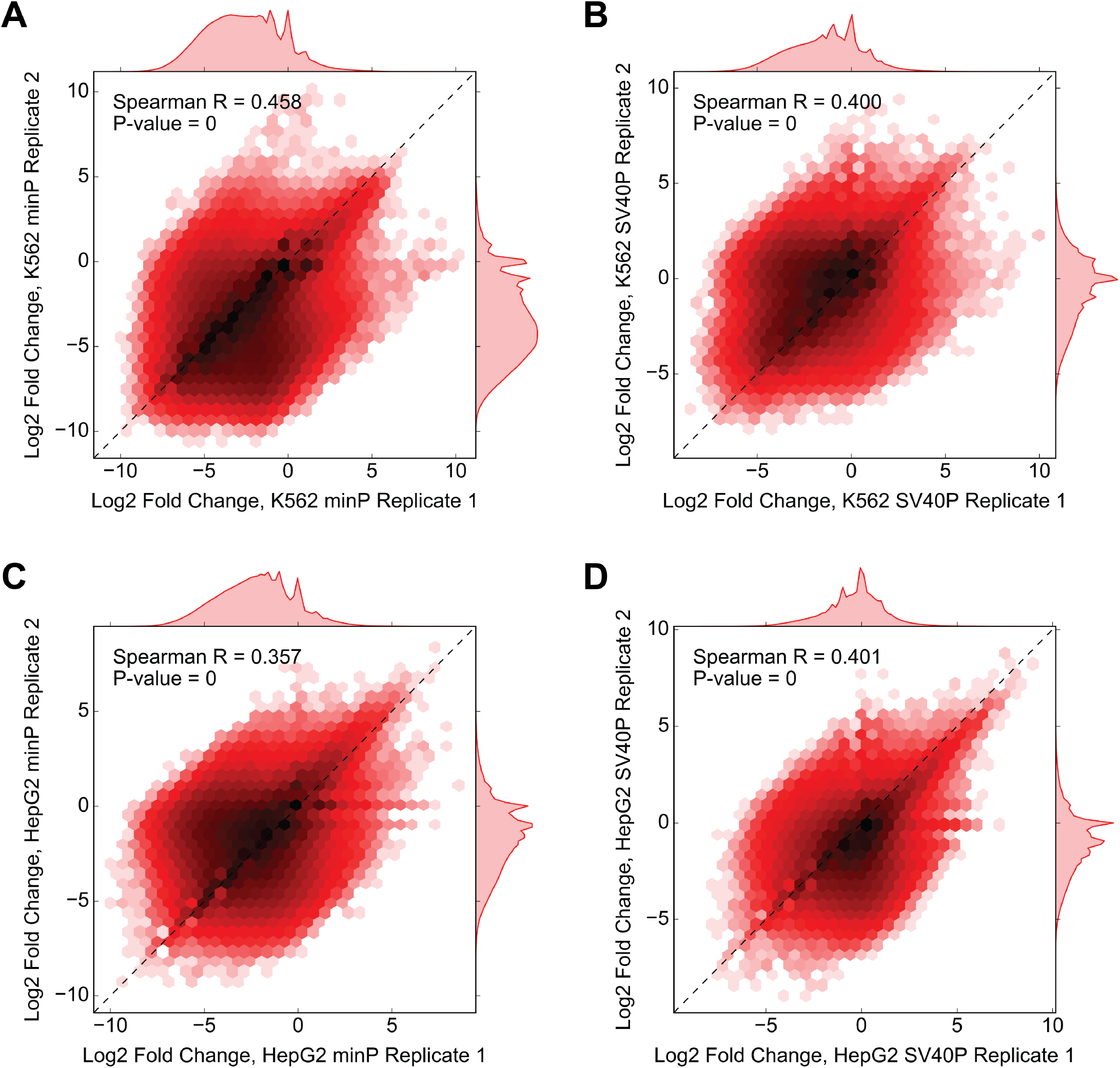
Between-replicate correlations for the four experimental groups in Sharpr-MPRA. (**A**) Correlation for fragments tested in K562 cells using the minimal promoter (minP); same plot as **Figure 1C** with added marginal distributions. *P*-values listed as 0 are less than Python’s float precision, i.e. *P* < 1E-300. (**B**) K562 cells with the SV40P promoter. (**C**) HepG2 cells with minP. (**D**) HepG2 cells with SV40P.

**Supplementary Figure 2:**
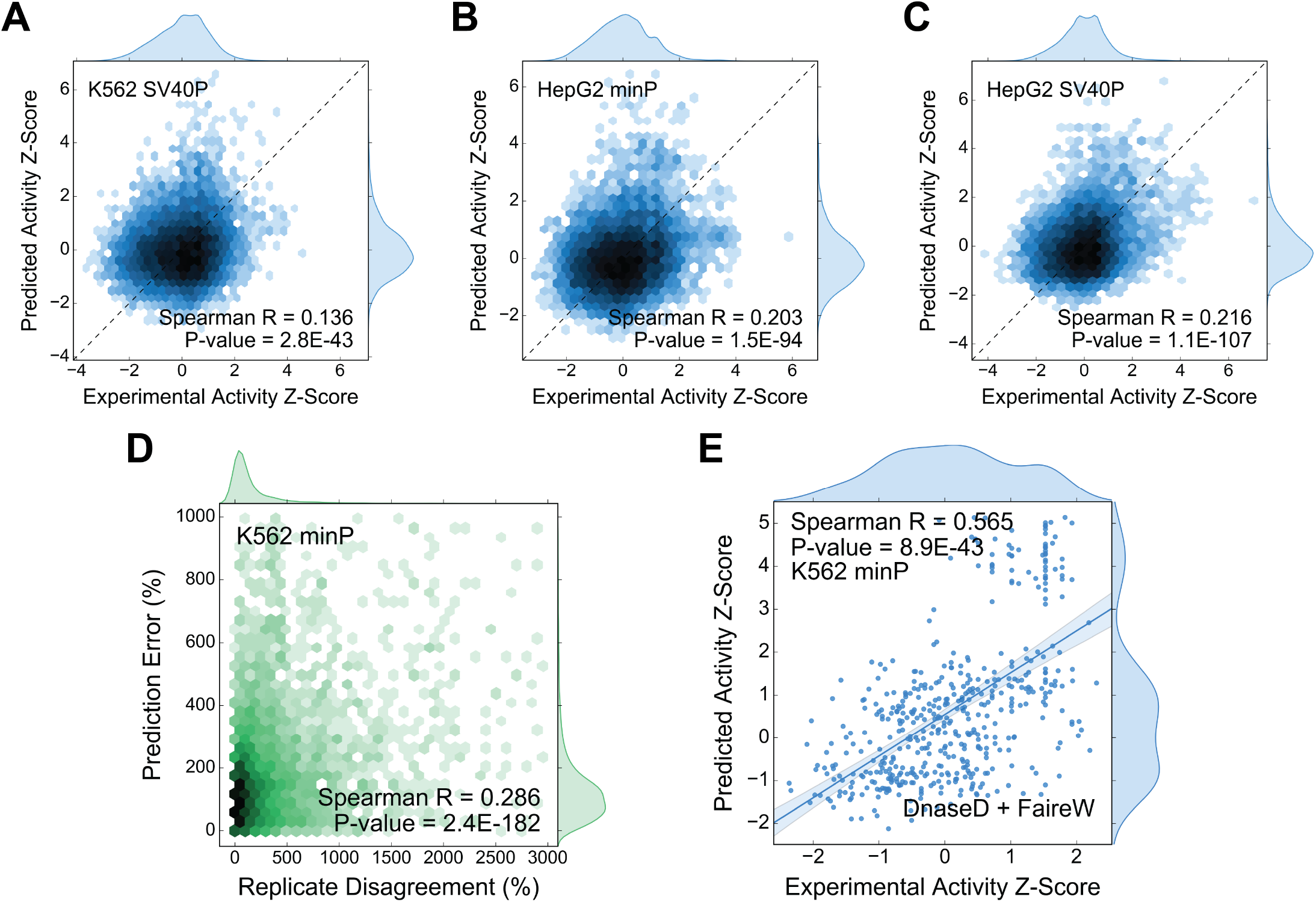
Detailed look at MPRA-DragoNN’s prediction performance. (**A**) Correlation between experimental regulatory activity *z*-scores and predicted regulatory activity *z*-scores for the K562 SV40P task (analogous to **Figure 2A**). These predictions are for fragments in the held-out test set (Sharpr fragments in chromosome 18). (**B**) Performance for HepG2 minP task. (**C**) Performance for HepG2 SV40P task. (**D**) Positive correlation between the difference in regulatory activity across replicates vs. prediction error, *i.e.*, fragments with more noisy experimental values have reduced prediction accuracy. (**E**) Improved prediction performance (*ρ* = 0.57) for fragments lying in accessible putative enhancers designated as ‘DnaseD’ or ‘FaireW’ states by ChromHMM annotations for the K562 cell type.

**Supplementary Figure 3:**
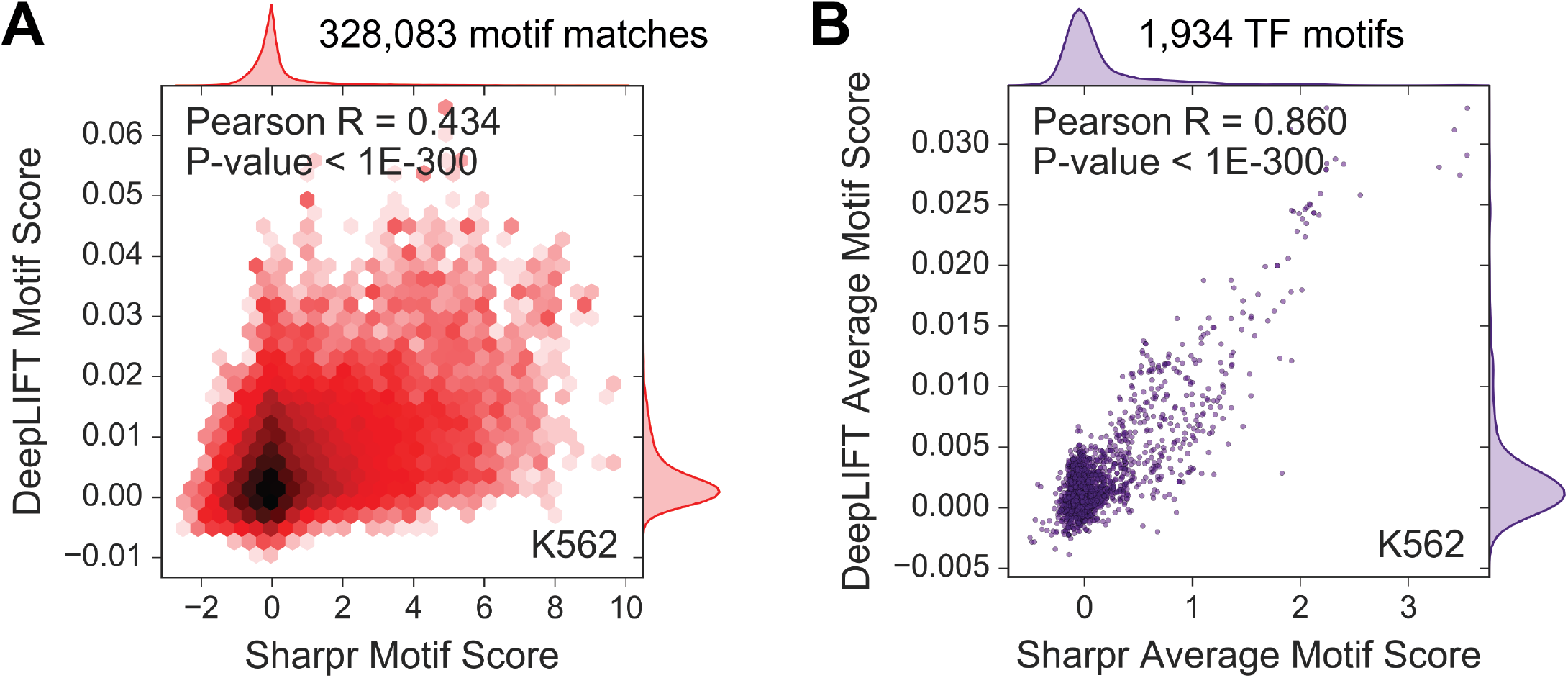
Agreement between DeepLIFT and SHARPR nucleotide scores at transcription factor motif matches. (**A**) Correlation between averaged DeepLIFT vs. averaged SHARPR nucleotide scores at each of the 328K motif matches that overlap at least one Sharpr-MPRA fragment. Each datapoint corresponds to a particular motif instance. (**B**) DeepLIFT vs. SHARPR correlation of the average motif match scores across all matches of one of the 1,934 different TF motifs. Each datapoint corresponds to a particular motif.

**Supplementary Figure 4:**
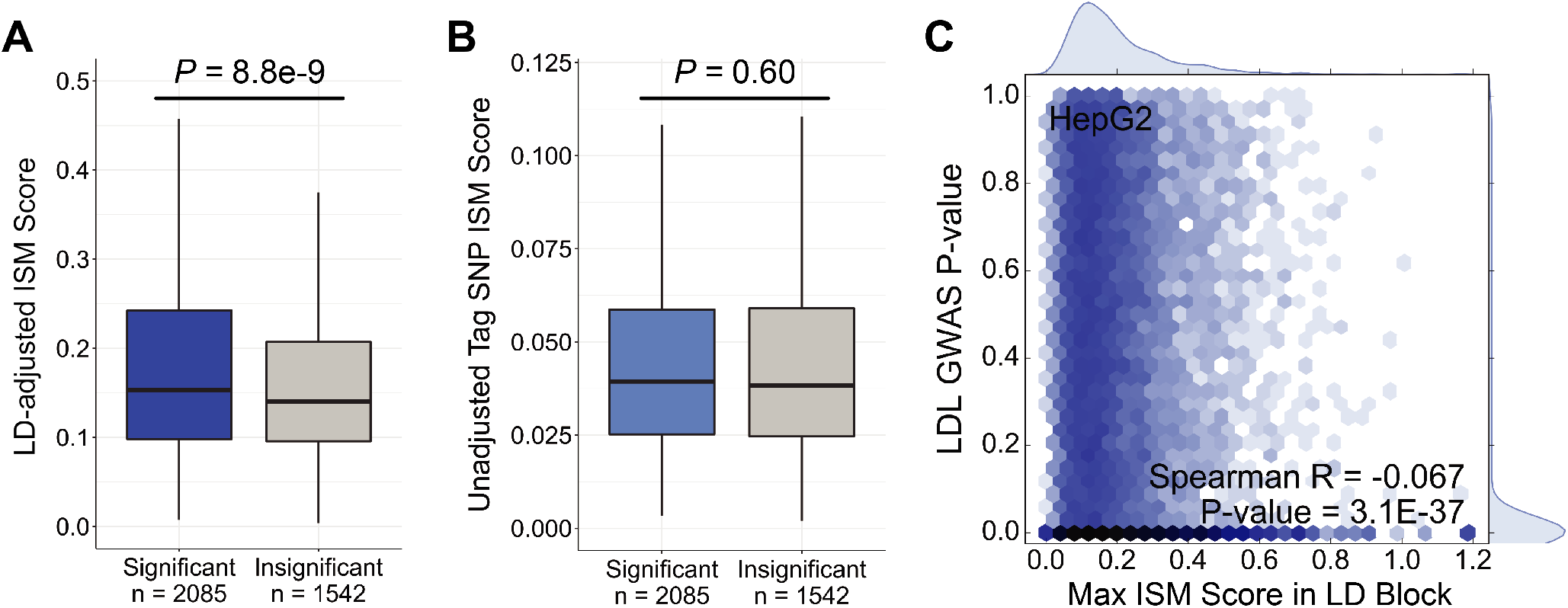
LD-adjusted ISM scores offer some discriminating power between significant and insignificant GWAS variants. (**A**) Boxplots showing distributions of LD-adjusted ISM scores (using the HepG2 minP model) for significant (*P* < 5×10^−8^) GWAS variants correlated with LDL cholesterol vs. insignificant (*P* > 0.1) GWAS variants. The significant variants are scored higher (*P* = 8.8 × 10^−9^). (**B**) Boxplots showing distributions of unadjusted ISM scores for significant vs. insignificant GWAS tag variants. Without accounting for linkage disequilibrium, there is no difference in scores between significant and insignificant variants. (**C**) Genome-wide correlation between LD-adjusted variant ISM scores and *P*-values of association of the variants with LDL cholesterol levels.

**Supplementary Figure 5:**
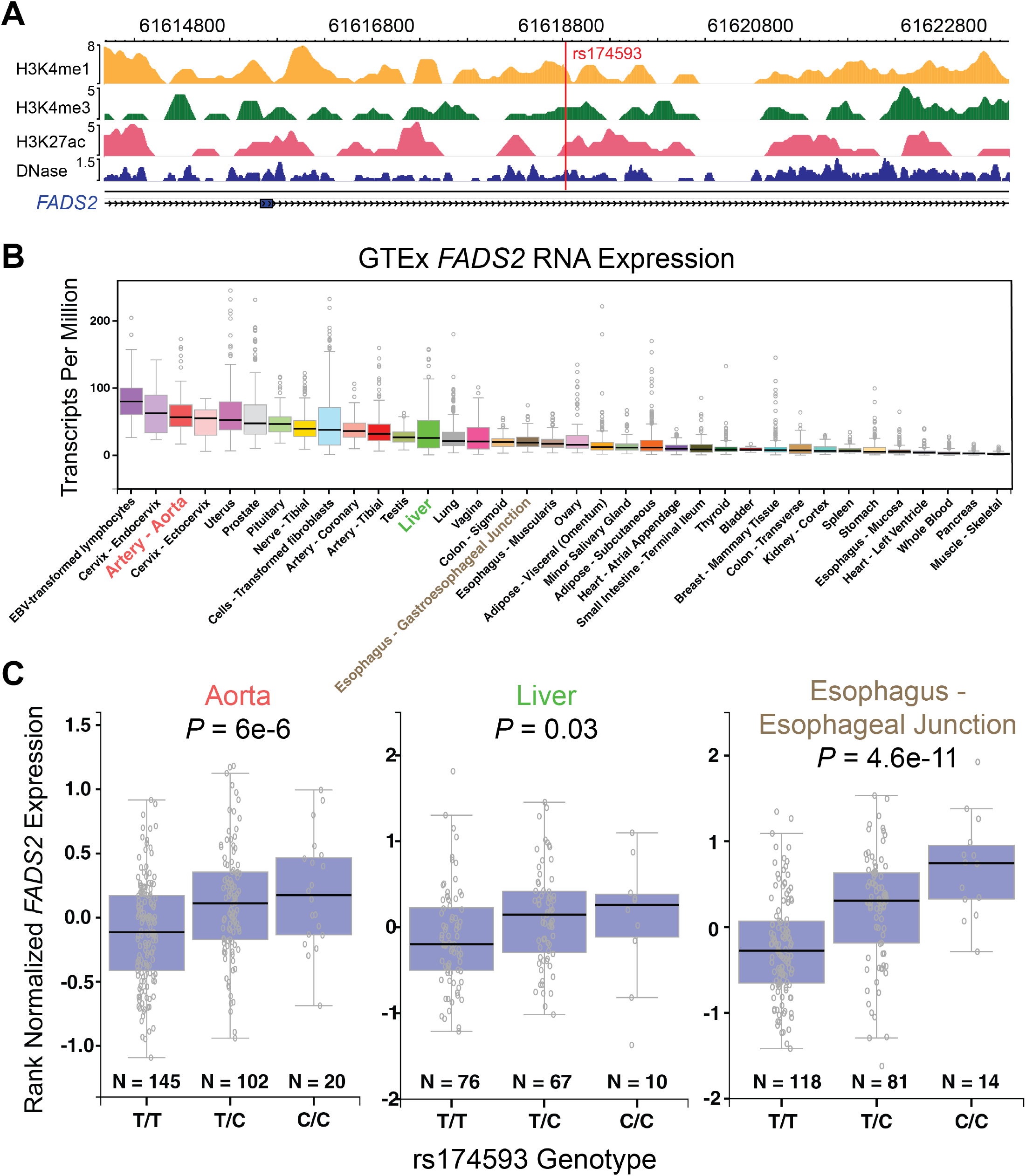
Further evidence for rs174593 as a regulator of FADS2 expression. (**A**) HepG2 epigenetic landscape in the genomic neighborhood of rs174593. (**B**) Sorted *FADS2* expression boxplots for a number of different tissue types from the GTEx consortium^60^. (**C**) *FADS2* expression vs. rs174593 genotype for three tissues that express *FADS2* from (B). Notably, aorta and esophagus cells display greater epigenomic activity at the rs174593 locus (data from Roadmap, not shown) and have stronger eQTL signal, suggesting that rs174593 may act as a disease variant through these (or other) tissues.

## Notes

#### Summary of Updates

The model was renamed from 'SNPpet' to 'MPRA-DragoNN'. Some sentences and statements in the text were revised for increased clarity and accuracy. The 'Availability' section was updated to point to a new Github repository with more usable software. Authors added.

https://github.com/kundajelab/MPRA-DragoNN/

